# A plug-and-play transepithelial/transendothelial electric resistance (TEER)-upgraded organ-on-chip system to measure barrier dynamics in real-time

**DOI:** 10.1101/2025.04.10.648170

**Authors:** Tim Kaden, Sophie Besser, Nader Abdo, Alexander S. Mosig, Knut Rennert, Sandor Nietzsche

## Abstract

The integration of transepithelial/transendothelial electrical resistance (TEER) measurement into organ-on-chip (OoC) platforms provides a unique opportunity to monitor the integrity of biological barriers in real-time. This is particularly important for detecting rapid changes in the temporal dynamics of intercellular junctional complexes in response to drug compounds, changes in host-microbiota interactions, and pathological disease states. Conventional TEER systems usually require special cell culture components or complex measurement technology that must be operated by experienced users. In this work, we present an innovative approach that represents an extension of the existing and well-established Dynamic42 chip platform by integrating semi-transparent TEER electrodes with fixed positions in combination with a measurement device. Remarkably, this system works in a plug-and-play manner and can continuously measure TEER inside the incubator without user intervention or invasive manipulation. Other important features of OoC, such as the microfluidic perfusion, multicellular cell colonization, and the possibility of microscopic examination, are not compromised by the integrated TEER electrodes. To demonstrate the performance of this new TEER system, we leveraged a 3D intestine-on-chip (IoC) model and investigated TEER during model assembly, barrier disruption, and recovery as a proof-of-concept. Moreover, we compared and discussed this with data from a conventional end-point fluorescence permeability assay to demonstrate the benefits of real-time measurements with higher sensitivity.

## 1. Introduction

The emergence of OoC technology represents a major step in biomedical research and provides a promising alternative to conventional *in vitro* and *in vivo* models. OoCs are microfluidic cell culture models that simulate the physiological functions of human organs and thus more accurately emulate the human biology [1–3]. They allow researchers to study complex cell-cell interactions in a controlled microenvironment, which is of great relevance for areas such as biological research, drug development, and personalized medicine.

A fundamental aspect of OoC technology is the implementation of compartmentalized tissue barriers under dynamic perfusion. The intestinal barrier is one example, which is involved in regulating the absorption of nutrients and drugs and, more importantly, tightly controls the spatial separation of luminal microbes from the intestinal epithelium, which also comprises cells of the intestinal immune system [4]. In addition to epithelial barriers, vascular barriers, such as the blood-brain-barrier (BBB), are critical to maintain a controlled exchange of molecules and substances between the blood circulation and adjacent organ tissues [5]. Evaluation of the integrity and functionality of these barriers is essential for investigating drug safety and distribution, inflammatory diseases, and microbial infections.

TEER measurement has been developed as a pivotal technique for assessing barrier integrity in OoC systems in real-time [6,7]. Compared to traditional *in vitro* methods, such as the use of “chopstick” type electrodes in Transwell cultures, TEER electrodes in chip platforms are incorporated at a fixed position without being movable and therefore provide a more accurate and stable measurement with less sensitivity distribution [8]. Moreover, in contrast to frequently used fluorescent tracer molecules as an endpoint assay to assess barrier permeability, TEER is a non-invasive, more sensitive, and real-time method to quantify barrier functionality over long periods of time [9]. Various platforms have successfully integrated TEER electrodes in different OoC models, for example in models of the intestine [6,7,10–13], blood-brain-barrier (BBB) [14–19], and lung [7,20]. While there are many established OoC models with TEER measurement, they still face certain limitations such as complicated fabrication, small molecule adsorption by polydimethylsiloxane (PDMS) as chip material, non-transparent electrodes, uneven current distribution, lack of scalability and reproducibility, and above all complicated user-unfriendly handling, which hinders widespread adoption [21].

Here, a non-PDMS monolithic biochip platform that has been upgraded with TEER electrodes and combined with an autonomously operating measurement device is described. The systems offer several innovative capabilities, including non-invasive real-time measurement for long-term experiments, an easy chip sterilization protocol, high reproducibility and sensitivity, simple handling, and scalability for different biological applications.

As a proof-of-concept, we leveraged an established IoC [22,23] model to validate barrier functionality with TEER measurement. The IoC model was probed with a barrier-disrupting chemical and observed for subsequent tissue recovery. Collectively, we demonstrated the potential of the system to explore tissue barrier dynamics during cell colonization, barrier breakdown, and tissue regeneration.

## 2. Method

### TEER-upgraded biochips

TEER-upgraded biochips (BC002T) were manufactured by Dynamic42 (Jena, Germany). Biochips were made from injection-molded polybutylene terephthalate (PBT) base bodies and consisted of two independent cavities for cell culture. Each cavity included a top and a bottom channel for cell seeding, which were separated by an integrated 12 µm thin polyester (PET) membrane (TRAKETCH Sabeu, Radeberg, Germany) with a pore diameter of 0.4 µm. The cavities were sealed with 0.125 mm thin polycarbonate (PC) foils, which were laser-welded on the top or bottom. The inner surfaces of the foils were equipped with electrodes. Their patterns were produced by sputtering with gold using a thickness that provides both electrical conductivity and optical transparency. The electrodes were connected via an electrical connector placed on the outside of the biochips.

### Electrode fabrication and integration

In order to integrate the electrodes into the biochip, a special design of the bonding foils was required to ensure a leak-proof connection with the read-out device.

The raw PC sheets were preprocessed to subsequently integrate the electrodes. First, the protective film was cut and selectively removed from the PC sheet at the regions of the electrodes. The top and bottom bonding foils were cut to the desired size for bonding the biochip base bodies. Electrodes were prepared by gold sputtering using Argon process gas at a vacuum pressure of 5 Pa (0.05 mbar), whereby a layer of 18 nm was deposited. Afterward, the foils were laser-welded onto the biochip base body using a laser welding device. The biochips were checked for quality and leakage after laser welding. Finally, a small, printed circuit board providing electrical connectors was attached to the biochip with integrated TEER electrodes.

### TEER read-out device

A newly designed handheld TEER measurement device was used (tighTEER T-31, Dynamic42 GmbH). It features an autonomous operation and is small enough to be placed next to the biochips in a cell culture incubator. It is battery-driven and simultaneously transmits resistance values of up to 6 different cavities via Bluetooth in real-time. Therefore, a generator module provides an alternating current (AC) voltage at a fixed frequency of 530 Hz with a fixed amplitude of 5 mV_RMS_. This voltage is distributed to 6 channels by means of an analogue multiplexer and the returning current is measured. After calculating the TEER resistance, a time stamp and all 6 data values are transmitted. The time interval for the measurements can be selected on a control panel between 5 sec and 60 min. Care has been taken to ensure long-term temperature stability of the analogue electronic circuitry by automatic calibration routines. A high-precision real-time clock provides accurate timing.

### Cell culture

All cultivation and incubation steps were performed in a humidified cell culture incubator at 37°C and 5% CO_2_, unless stated otherwise. Cells used in this study were tested negative for mycoplasma contamination using the Venor^®^GeM Classic Kit (Minerva Biolabs, Berlin, Germany).

HUVECs were isolated under ethical approval 20-20-1684, 3939-12/13 following written informed consent from donors.

Cryopreserved HUVECs were thawed and seeded in culture flasks at a density of 1.3 × 10^4^ cells/cm^2^. The cells were cultured in Endothelial Cell Growth Medium (ECGM) MV (Promocell, Heidelberg, Germany) with supplements and 1× antibiotic antimycotic solution (AAS, Merck, Darmstadt, Germany). Medium was exchanged every two days until further passaging or seeding into biochips. HUVECs were used up to passage 3.

Caco-2 epithelial cells (acCELLerate, Hamburg, Germany) were thawed and seeded in culture flasks at a density of 0.8 × 10^4^ cells/cm^2^. Cells were cultured in gut maintenance medium (GMM) containing DMEM with 4.5 g/L glucose (Lonza, Cologne, Germany), 10% fetal bovine serum (FBS, Capricorn Scientific, Ebsdorfergrund, Germany), 1 mM sodium pyruvate solution (Capricorn), 1× MEM non-essential amino acids (Capricorn), 5 mg/mL holo-transferrin (Merck) and 20 µg/mL gentamycin (Merck). Medium was exchanged every three to four days until further passaging or seeding into biochips.

Human monocyte-derived macrophages (MDMs) were differentiated from human peripheral blood mononuclear cells (PBMCs), which were isolated from blood of healthy volunteers. All donors were informed about the study and gave written consent. All procedures were performed according to the approved guidelines and regulations and to the guidelines set forth in the Declaration of Helsinki. The isolation of PBMCs and pre-differentiation to MDMs were performed as previously described [23,24]. Briefly, PBMC were isolated from blood by density gradient centrifugation. Adherent monocytes were cultured in X–VIVO 15 (Lonza) supplemented with 10% human autologous serum and 1× AAS. Differentiation into MDMs was induced by adding 10 ng/mL macrophage colony-stimulating factor (M-CSF, PeproTech, Hamburg, Germany) and 10 ng/mL granulocyte-macrophage colony-stimulating factor (GM-CSF, PeproTech) to the culture medium for at least five days.

### Intestinal model assembly

Intestinal models were gradually assembled as previously described [22,23]. Briefly, biochips were sterilized, washed, and subsequently coated in the top channel with 50 µg/mL bovine collagen A (PAN-Biotech, Aidenbach, Germany) for 1 h at 37°C and 5% CO_2_. HUVECs were seeded in the top channel on collagen-coated membranes at a density of 1.38 × 10^5^ cells/cm^2^ (in total 3 × 10^5^ cells) in supplemented ECGM MV with AAS. Medium was exchanged daily under static conditions for 72 h. Prior to the seeding of Caco-2 in the opposite lower channel, the membrane side facing the bottom channel was coated with 50 µg/mL bovine collagen A for 1 h at 37°C and 5% CO_2_. Coating of this membrane side was achieved by incubating the biochips upside down with closed inlets and outlets. Caco-2 cells were seeded in the bottom channel at a density of 2.16 × 10^5^ cells/cm^2^ (in total 3.5 × 10^5^ cells) in gut seeding medium (GSM) containing GMM with 20% FBS. Biochips were statically incubated upside down for 24 h. Intestinal models were perfused by a peristaltic pump system (Masterflex, VWR International, Bruchsal, Germany) in both channels at 50 µL/min equaling shear stress rates of 0.013 dyn/cm^2^ (0.0013 Pa) in the top channel and 0.006 dyn/cm^2^ (0.0006 Pa) in the bottom channel.

### TEER measurement and treatment

After establishing the 3D IoC model, the tighTEER T31-device (Dynamic42 GmbH) was used to monitor the integrity of the cell barrier. The device itself can be placed in the incubator with the biochips and requires no user intervention. Wirelessly connected via Bluetooth, the status of the cell barrier can be measured and logged in real-time on an external computer. The transmission protocol is based on a virtual communication (COM) port and transmits time stamps and resistance values in plain text. In our experiments we used 1 min or 20 min intervals and the open source software Better Serial Plotter (https://github.com/nathandunk/BetterSerialPlotter, version v0.1.2), which is a drop-in replacement for the Arduino serial plotter (https://www.arduino.cc/) software.

Models were equilibrated for 30 min before the treatment. Ethylenediaminetetraacetic acid (EDTA, Merck) was added at a concentration of 10 mM into the perfusion reservoirs of the intestinal channel to induce barrier disruption. Treatment with EDTA was performed for 30 min. EDTA was replaced by fresh GMM in the intestinal perfusion reservoir to initiate tissue recovery over 24 h. Medium was also exchanged in control models to exclude TEER changes in response to ion and temperature fluctuations. TEER measurement was stopped after recovery.

The resistance of the cell barrier R_Cells_ was calculated as previously described [8,25]:

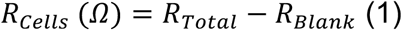

in which R_Total_ is the resistance across the cell layers and the porous membrane and R_Blank_ is the resistance without the cell layers.

Based on this equation, the TEER resistance R_TEER_ on a defined barrier area A can be calculated using the following formula:

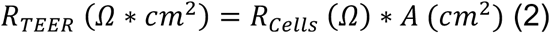

### Permeability assay

Intestinal permeability was assessed by diffusion of fluorescein isothiocyanate– dextran (FITC-dextran, 3-5 kDa, CAS number: 60842-46-8, Merck). Medium in both channels was replaced by prewarmed phenol-red free William’s medium E (PAN-Biotech). FITC-dextran was diluted in phenol-red free William’s medium E to a stock concentration of 1 mg/mL. A volume of 250 µL of the stock solution was transferred twice in the intestinal bottom channel to minimize dilution effects. All inlets and outlets of the biochip were plugged, biochips were turned upside down and incubated in the dark for 1 h at 37°C and 5% CO_2_. After incubation, supernatants were collected for both top and bottom channels and transferred into a black 96-well microplate (Corning, Amsterdam, The Netherlands). Fluorescence was measured in a microplate reader (INFINITE 200 PRO, Tecan, Crailsheim, Germany) at 492 nm (excitation)/ 518 nm (emission). FITC-dextran concentrations were calculated based on the plotted standard curve of each assay. The apparent permeability coefficient was calculated from permeated FITC-dextran concentrations using the following equation [26]:

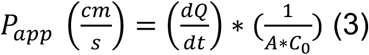

where dQ/dt is the steady-state flux within the incubation time (μg/s), A is the surface area of cultivated cells on the membrane (cm^2^), and C_0_ is the initial concentration of the FITC-dextran stock solution (μg/cm^3^).

### Cell viability assay

Cell viability was measured in chip-based models with and without integrated TEER electrodes using the CellTiter-Glo Luminescent Cell Viability Assay (Promega, Walldorf, Germany). Excised membranes from biochips were transferred to a microplate containing phenol-red free William’s medium E. The same volume of CellTiter-Glo Reagent was added (1:1 ratio). The assay was performed as described in the manual provided by the manufacturer. Luminescence was measured in a microplate reader (INFINITE 200 PRO) with an integration time of 1 s. ATP concentrations were calculated based on the plotted ATP standard curve of each assay.

### Immunofluorescence staining

Both channels were washed with cold PBS (4°C) with calcium and magnesium (PBS +/+) and adherent cells were fixed with ice-cold (−20°C) methanol (Carl Roth, Karlsruhe, Germany) for 15 min at -20°C. Channels were washed three times with PBS +/+ to remove the methanol. Membranes with fixed cells were recovered from the biochips by excision with a sharp scalpel. Membranes were transferred into permeabilization/blocking solution containing PBS +/+, 0.1% saponin (Carl Roth), and 3% normal donkey serum (Abcam, Amsterdam, The Netherlands) and were incubated for 30 min at RT. Afterward, membranes were halved and incubated in permeabilization/blocking solution containing the following primary antibodies: mouse anti-CD31 (1.66 µg/mL, Cell Signaling Technology, Leiden, The Netherlands), rabbit anti-von Willebrand Factor (vWF, 9.3 µg/mL, Agilent, Waldbronn, Germany) for vascular cell layers and mouse anti-E-Cadherin (2.5 µg/mL BD Bioscience, Heidelberg, Germany), rabbit anti-zonula occludens 1 (ZO-1, 1.25 µg/mL Thermo Fisher Scientific, Schwerte, Germany) for intestinal cell layers at 4°C overnight. Membranes were washed with PBS +/+ and 0.1% saponin and were incubated with the following secondary antibodies: DAPI, donkey-anti-mouse-AF555, and donkey-anti-rabbit-AF488 (all 10 µg/mL, Thermo Fisher Scientific) for 1 h at RT. Subsequently, membranes were washed and embedded in fluorescence mounting medium (Agilent, Glostrup, Denmark) between two microscopic glass slides.

### Image acquisition and analysis

Light microscopic images were taken to monitor tissue integrity during model assembly and TEER measurements. Images were acquired using the ZEISS Primovert phase contrast inverted cell culture microscope (Carl Zeiss AG, Jena, Germany) equipped with a Plan-Achromat 4x/0.1 or 20x/0.3 objective.

Fluorescence images were acquired using the ZEISS Axio Observer 7 (Carl Zeiss AG) with Apotome 3 and Plan Apochromat 20x/0.8 M27 objective. The ZEISS ZEN 3.7 software (Carl Zeiss AG) was utilized for microscope controlling, image acquisition, and processing. Images were obtained as Z-stacks with an interval of 1 µm (vascular) or 2 µm (intestinal) and were processed to maximum intensity projections after ApoTome Raw Convert.

### Statistical analysis

Statistical analysis was performed using GraphPad Prism v10.4.1 (GraphPad Software, La Jolla, CA, USA). The statistical tests and multiple comparison tests are noted in the respective figure legends. A p-value < 0.05 was considered statistically significant.

## 3. Results

Integrating non-invasive TEER electrodes at a fixed position (**figure 1(a)**) is a major advantage compared to traditional measurements using chopsticks. The electrodes were connected by an interface on the attached circuit board to the measurement device (**figure 1(b)**). The TEER values were recorded by the tighTEER T-31 device and wirelessly transferred to a computer, which was placed outside the cell culture incubator for real-time measurement without user interference (**figure 1(c)**). The IoC model for barrier disruption and recovery experiments consisted of a vascular cell layer formed by HUVECs in the top channel and an intestinal cell layer of Caco-2 cells in the bottom channel (**figure 2(a)**). Both cell layers were separated by a porous membrane, which enabled cell-cell communication and molecule diffusion but restricted cell migration due to the small pore size of 0.4 µm. Based on the transparency of the TEER electrodes, it was possible to continuously monitor cell attachment and growth with brightfield microscopy (**supplementary figure 1**). IoC models were assembled under static conditions (**supplementary figure 2**, **figure 2(c)**). After seeding HUVECs and Caco-2 cells, the models were connected to peristaltic perfusion (**figure 2(d)**). The models were pre-perfused for six days prior to experimental barrier disruption and recovery (**supplementary figure 2**). TEER values were recorded during static model assembly and perfusion by connecting the tighTEER T-31 device to the biochip (**figure 2(d)**). Upon seeding HUVECs, the TEER values slightly increased over 48 h before seeding the MDMs on top (**figure 2(e)**). MDMs were included in this experiment as an immune cell component to investigate their influence on barrier integrity. Seeding of MDMs led to a small elevation in TEER compared to cavities that were only lined with HUVECs. However, during seeding and formation of the confluent endothelial cell layer, TEER values remained low, with values < 50 Ω*cm^2^. A significant increase in TEER was observed after seeding Caco-2 cells in the bottom channel. The Caco-2 cells formed a tight barrier over 24 h, which resulted in TEER values > 200 Ω*cm^2^. Connecting the models to perfusion in both channels initially increased TEER values to approximately 250 Ω*cm^2^ and remained stable > 200 Ω*cm^2^ over 24 h of perfusion. Models incorporating only HUVECs or HUVECs with MDMs were also connected to the perfusion, although TEER values only minorly increased over the perfusion period (**figure 2(e)**). Besides, determination of cell viability in IoC models without or with integrated TEER electrodes did not reveal effects of the electrodes on cell viability, validating the biocompatibility of the electrodes (**figure 2(f)**).

**Figure 1:**
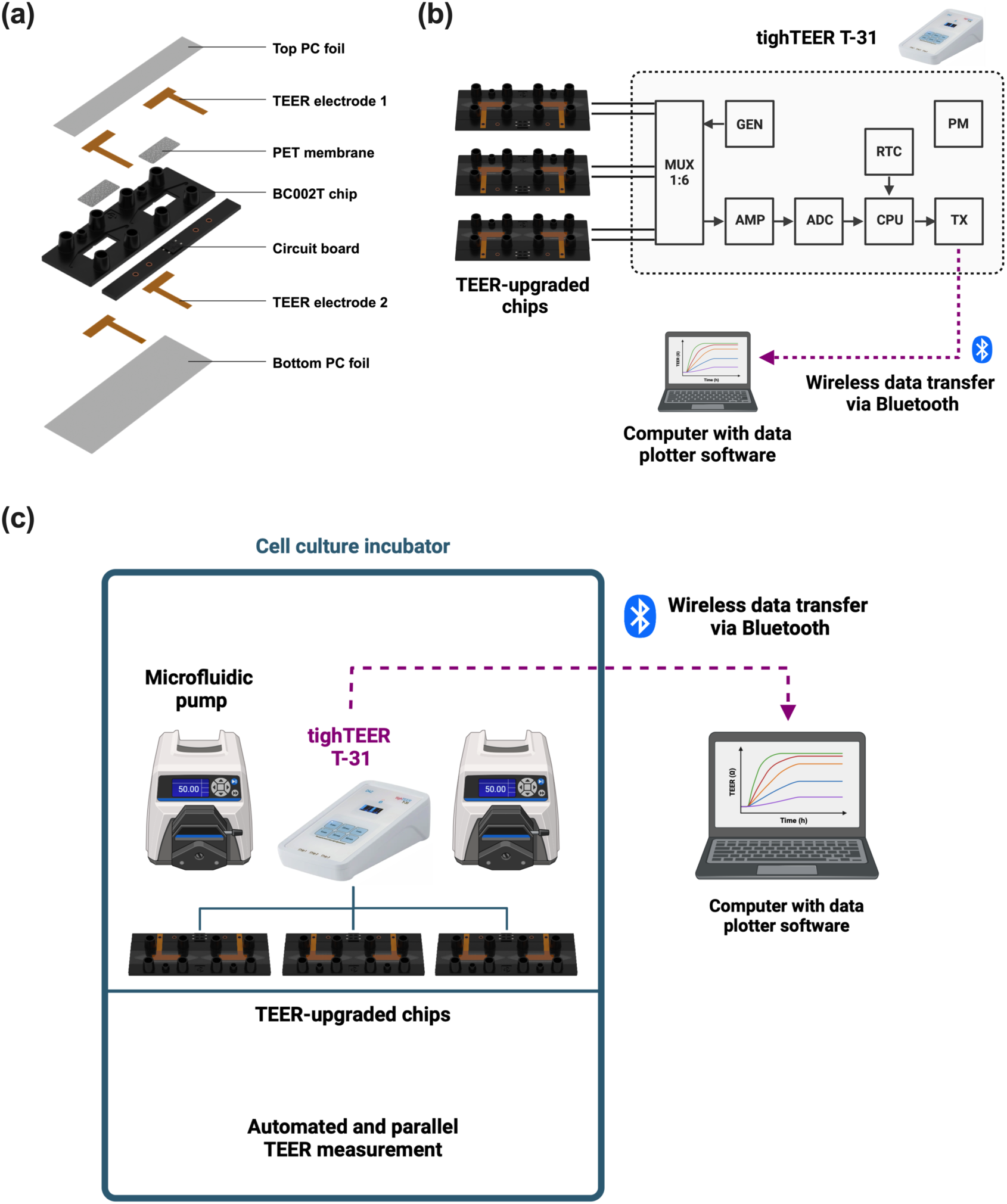
Integration of TEER electrodes into the Dynamic42 biochip platform and combination with the tighTEER T-31 measurement device. **(a)** Exploded view of the TEER-upgraded Dynamic42 biochip. **(b)** Schematic representation of the measurement setup. All of the following modules are integrated in an easy-to-use handheld device: sinewave voltage generator (GEN), 6-fold multiplexer (MUX) to support 3 biochips with 2 cavities each, signal amplifier (AMP), analog-to-digital converter (ADC), processing unit (CPU) managing the timing schedule and data calculation, real-time clock (RTC), wireless transmission unit (TX), and a power module (PM) for battery management. **(c)** Schematic illustration of TEER measurement setup. TEER-upgraded chips are cultured in a cell culture incubator under static or perfused conditions. Chips are connected via the attached circuit board to the tighTEER T-31 measurement device, which transfers the data via Bluetooth to an external computer with data plotter software. All figures were created with BioRender.com.

**Figure 2:**
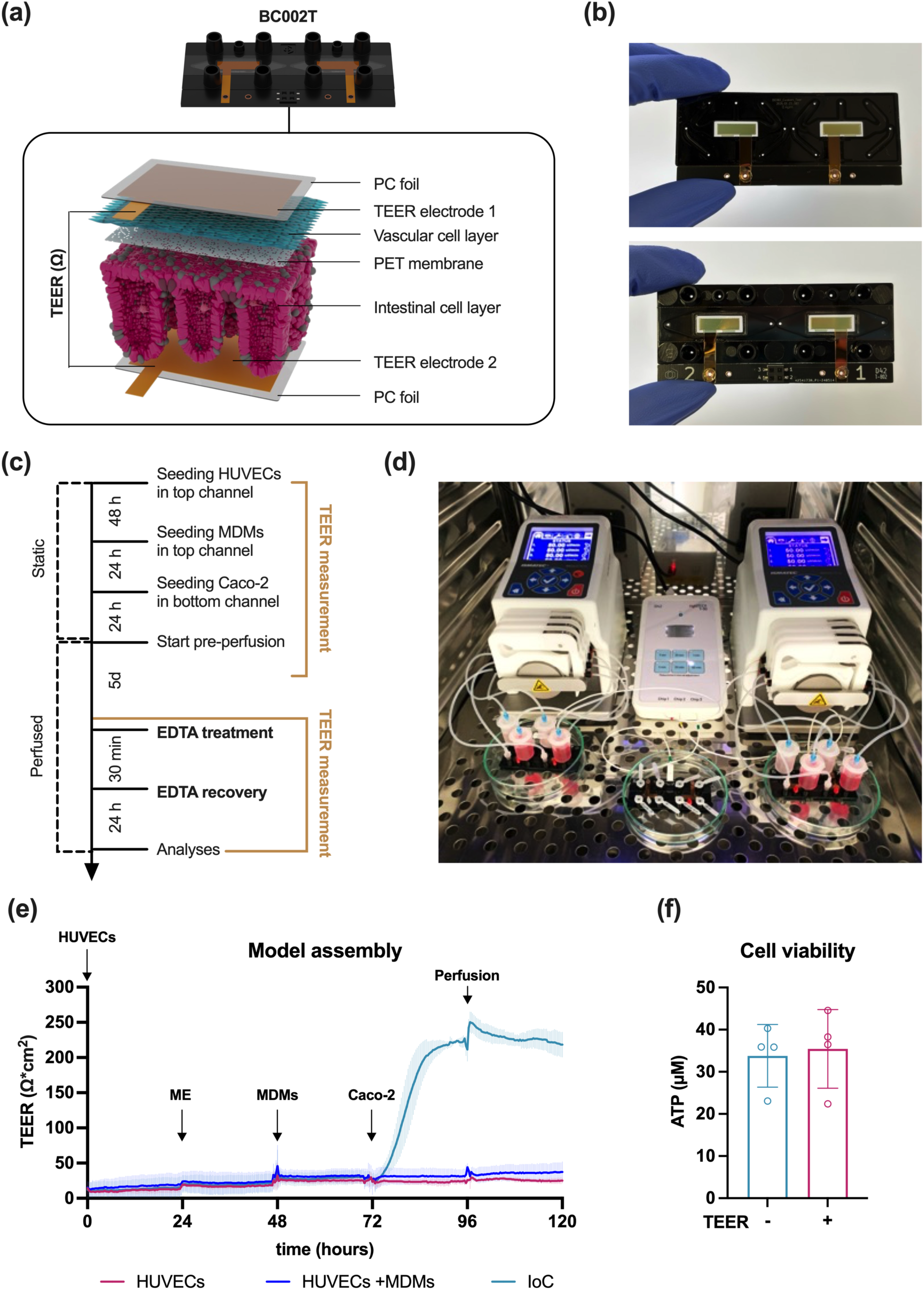
Outline of the IoC model with integrated TEER electrodes. **(a)** Enlarged section of the IoC model in the Dynamic42 biochip platform with integrated TEER electrodes. The model consists of a top channel with HUVECs (turquoise) forming the vascular cell layer and a bottom channel with Caco-2 cells (purple) forming the intestinal cell layer. Gold electrodes are placed in the chip on the inside of the bottom and the top bonding foil facing the cell layers. **(b)** Representation of the Dynamic42 biochip with integrated gold electrodes and attached circuit board with interface for the tighTEER T-31 device. Upper image: bottom side of the chip. Bottom image: top side of the chip. (**c)** Timeline of static intestinal model assembly, subsequent perfusion, and TEER measurement during model assembly or EDTA treatment and recovery. **(d)** Experimental model setup in a cell culture incubator. The tighTEER T-31 device is connected to the interface on the circuit board attached to the biochips (white cables). **(e)** TEER measurement during static assembly of the IoC model and subsequent connection to the perfusion. Measurements were done in chips with HUVECs (magenta), co-culture of HUVECs and MDMs (blue), or complete IoC models consisting of HUVECs, MDMs, and Caco-2 (cyan). TEER values were measured every 20 min and calculated using eq. (2) and the culture area of 1.32 cm^2^. Perfusion was initiated after 96 h of static assembly. ME = medium exchange. Data points represent mean connected by lines ± SD of 3 independent experiments (n = 3). **(f)** Measurement of cell viability in intestinal models without (-) or with (+) integrated TEER electrodes. Viability was measured as ATP concentration (µM). Bars represent mean ± SD of 4 independent experiments (n = 4). No statistical significance was found using a two-tailed paired t test.

As a proof-of-concept, the barrier integrity in the IoC model was manipulated and continuously monitored by TEER measurements. Ethylenediaminetetraacetic acid (EDTA), a strong metal ion chelator, was administered in the bottom chip channel at the intestinal epithelial barrier to induce barrier leakage (**figure 3(a)**). EDTA is known to bind divalent ions, like magnesium and calcium. The latter is important to regulate and maintain the cellular junctional complex, including adherens and tight junctions [27,28]. Light microscopic monitoring of the intestinal epithelial cells revealed cell rounding and detachment from the membrane after 30 min of EDTA treatment (**figure 3(b)**). In contrast, replacement of EDTA medium with fresh medium without EDTA induced tissue recovery within 24 h. The tissue recovered to the 3D morphology with villus- and crypt-like structures as it was observed for control models prior to EDTA treatment. This indicates that EDTA induced barrier breakdown rather than inducing cytotoxicity. Before introducing EDTA into the system, TEER values of the models were monitored during pre-perfusion before starting experimental manipulations. It was observed that TEER values were stable over 140 h of pre-perfusion with low variability across three biological replicates (**figure 3(c)**). As observed during the model assembly (**figure 2(e)**), medium exchange always resulted in spikes in TEER values, with moderate variability due to temperature and ion fluctuations. In addition, between the first and the second medium exchange was a gap of 72 h associated with a slight decrease in TEER. This might be due to medium consumption over time but could also be a result from the formation of 3D villus- and crypt-like structures in the IoC reflecting a remodeling of the tissue and, most importantly, cellular junctions. When models were treated with 10 mM EDTA after 30 min equilibration, TEER values significantly decreased and reached almost baseline levels (**figure 3(d)**). The decrease in TEER was rapidly induced after treatment with EDTA, whereas control-treated models maintained stable TEER levels of approximately 150 Ω*cm^2^. Upon recovery, TEER values increased to control levels within 24 h, suggesting a complete restoration of the barrier functionality comparable to levels prior to EDTA treatment (**figure 3(e)**). While the standard deviation in these experiments reflects the biological variability, TEER values from barrier disruption and recovery experiments were blank corrected, normalized, and individually plotted to showcase also the high measurement accuracy of the tighTEER T-31 device (**supplementary figure S3(a), (b)**).

**Figure 3:**
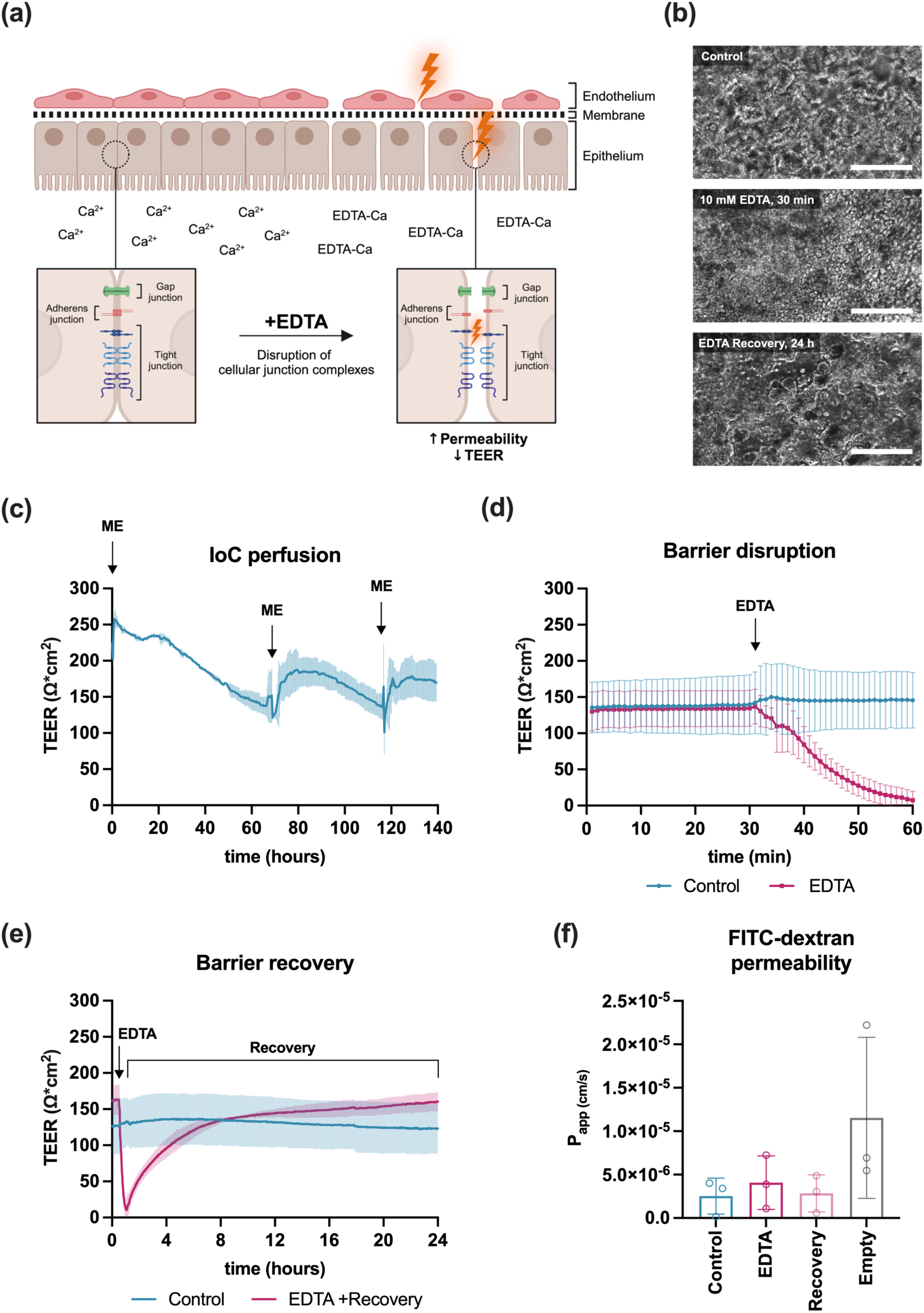
Monitoring of barrier disruption with TEER measurement. **(a)** Schematic representation of EDTA-induced barrier breakdown in the IoC model. **(b)** Light microscopic monitoring of intestinal morphology in perfused control models (top image), EDTA-treated models (30 min, middle image), and recovered models after EDTA cessation (24 h, bottom image). Scale bars: 200 µm. **(c)** Monitoring of TEER (in Ω*cm^2^) during perfusion of the IoC model. ME = medium exchange. TEER values are plotted as mean connected by lines ± SD of 3 independent experiments (n = 3). **(d)** Measurement of TEER during chemically induced barrier disruption. TEER (in Ω*cm^2^) was measured every minute over a total period of 1 h. EDTA (10 mM, black arrow) was administered after 30 min pre-equilibrium. Values are plotted as mean connected by lines ± SD of 3 independent experiments (n = 3). **(e)** Induction of barrier recovery after EDTA cessation. TEER values (in Ω*cm^2^) were measured every minute over a recovery period of 24 h. TEER values are plotted as mean connected by lines ± SD of 3 independent experiments (n = 3). **(f)** Permeability of FITC-dextran in IoC models after EDTA treatment for 30 min or recovery over 24 h. Empty = empty chip with 0.4 µm pores. Permeability is shown as the calculated apparent permeability coefficient (Papp). Bars present mean ± SD of 3 independent experiments (n = 3). No statistical significance was found using a one-way ANOVA with Tukey’s multiple comparisons test. **Figure 3(a)** was created with BioRender.com.

In comparison, measurement of barrier permeability with a conventional FITC-dextran permeability assay revealed only slight differences between the test conditions, highlighting the low sensitivity of this assay in determining barrier alterations (**figure 3(f)**). While a minor trend of increase was measured for EDTA treatment and the induction of the recovery slightly reduced this, there were no significant differences. The highest permeability was measured in empty chips without cells and 0.4 µm pores. While EDTA treatment induced a small increase in permeability, the differences to control models are very small, while TEER measurements (**figure 3(d)**) clearly indicated significant changes in a real-time manner.

To validate underlying changes in cellular junctions upon barrier dysfunction and recovery, immunofluorescence staining of both intestinal epithelial cells and vascular endothelial cells was performed (**figure 4**). Here, it was shown that untreated control models expressed defined networks of the adherens junction protein E-cadherin and the tight junction protein ZO-1 in the 3D intestinal cell layer. Furthermore, the presence of CD31, an adhesion molecule that is enriched at endothelial intercellular junctions [29], and the typical endothelial cell marker vWF were shown to be expressed in the vascular cell layer. Both intestinal and vascular junction proteins were disrupted after treatment with 10 mM EDTA for 30 min. The breakdown of intestinal junction complexes was further accompanied by a depletion of fluorescence signals from cell boundaries and increased signal distribution in the cytoplasm. Despite a reduction in junctional markers, both cell layers showed a homogenous distribution of DAPI-positive nuclei across the images, indicating that EDTA treatment for 30 min is affecting junctional complexes, but not nuclei integrity. After 24 h of recovery, both cell layers demonstrated a restoration of the junctional proteins E-cadherin, ZO-1, and CD31 as it was observed for untreated models.

**Figure 4:**
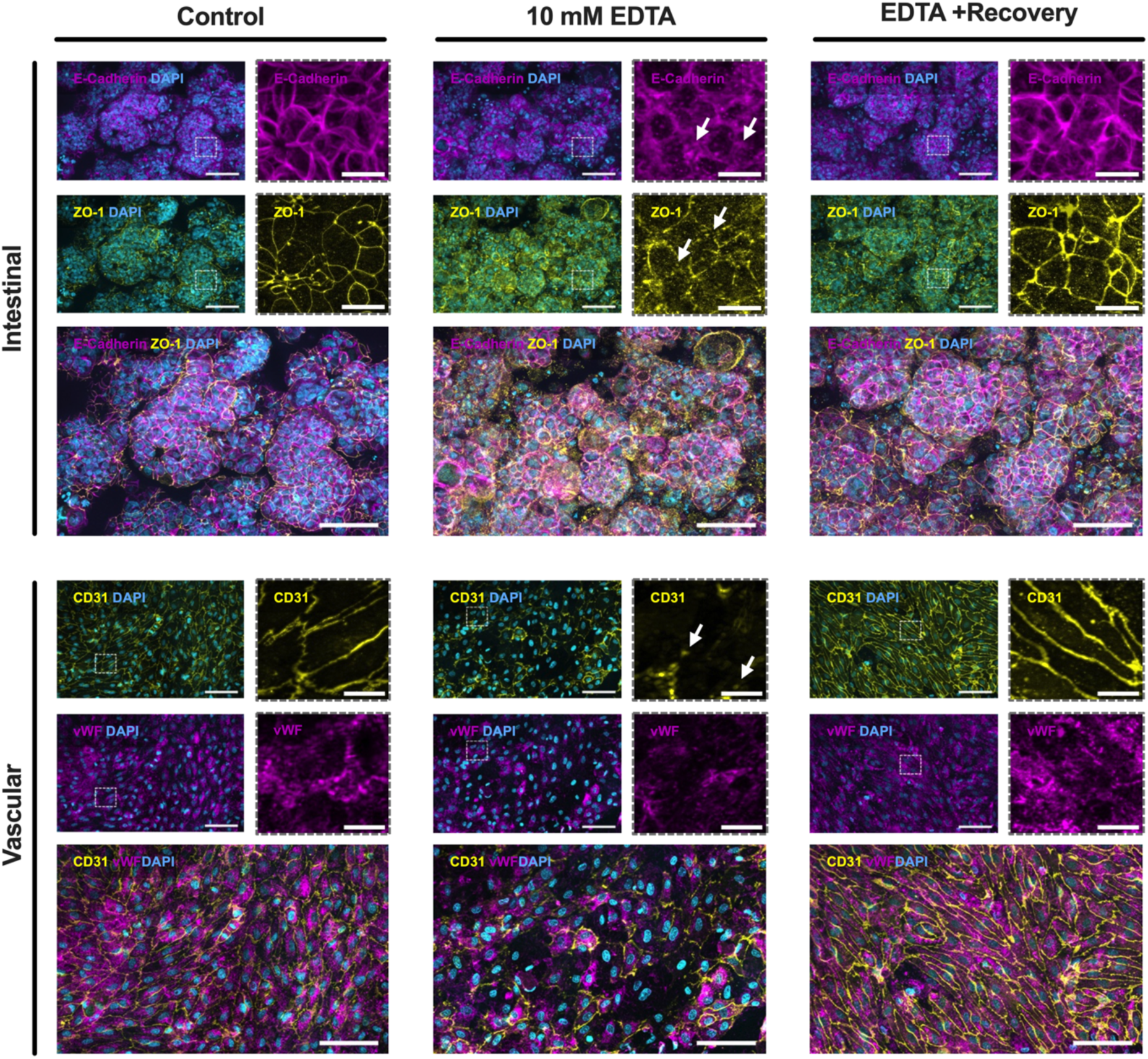
Assessment of junction complex disruption in the IoC model. Representative immunofluorescence images of intestinal and vascular cell layers in the IoC model after TEER measurements. Intestinal cells were stained for the adherens junction protein E-Cadherin (magenta), tight junction protein ZO-1 (yellow), and nuclei (DAPI, blue). Vascular cell layers were stained for CD31 (yellow), von Willebrand factor (vWF, magenta), and nuclei (DAPI, blue). White arrows indicate disruption of junctional complexes or membrane-associated adhesion molecules. Scale bars: 100 µm for uncropped sections (with DAPI), 20 µm for enlarged image sections (dotted line, without DAPI).

## 4. Discussion

In this work, we designed and integrated fixed semi-transparent gold electrodes into a monolithic non-PDMS biochip to measure TEER across tissue barriers. By combining the TEER chip with the tighTEER T-31 measurement device in a simple plug-and-play manner, barrier integrity could be measured at various intervals in real-time without user interference in a cell culture incubator. This enabled us to measure TEER up to eleven days, although this time limit was only due to the experimental setup and could otherwise be extended. By integrating electrodes that cover almost the whole membrane area for tissue culture, it allowed for the quantification of normalized TEER in Ω*cm^2^, as it was previously proposed [25,30–32], and permitted a comparison to similar chip models or Transwell systems. This is in particular important to extrapolate and compare data generated in systems with different chip geometries and electrode designs.

Herein, stable and reproducible data were collected during static and perfused IoC model assembly and subsequent manipulation of the intestinal barrier by EDTA. We were able to demonstrate increasing TEER values in correlation with higher complexity during the model assembly. HUVECs or the co-culture of HUVECs with MDMs exhibited relatively low levels of resistance between 10 and 40 Ω*cm^2^, which is in line with previous measurement of TEER in a BBB-chip model consisting of endothelial cells [33]. In contrast, adding epithelial Caco-2 cells in our model significantly increased TEER due to their rapid formation of a tightly connected monolayer and increasing barrier function during differentiation [34]. Upon perfusion, a slight decrease of TEER was noted when there was no medium exchange for two to three days. It is assumed that the culture medium is altered by the cells, resulting in depletion of tight junction-stabilizing ions, induction of pH changes, or secretion of metabolic products. On the other hand, TEER values consistently increased directly after exchanging the medium. In general, we measured very consistent data across the biological replicates, with moderate deviations during medium exchanges.

Upon perfusion, values between 100 and 200 Ω*cm^2^ were measured in the IoC model. This is similar to values that were observed in other OoC models integrating intestinal epithelial cells. For example, Lucchetti et al. showed variable TEER values along a chip channel, reaching a maximum of 125 Ω*cm^2^ [12]. Besides, they highlighted chemical gradients, temperature, pressure, and air bubbles as potential factors to influence TEER values across the culture area, which was the rationale in their study to measure TEER at four different positions along the channel. In addition, another study measured TEER in a co-culture of HT29 tumor spheroids with MDCK_VB6-CB cells, showing values between 50 and 80 Ω*cm^2^ [35]. A recent publication outlining a bioprinted hydrogel gut-on-chip model reported values up to 80 Ω*cm^2^ in a co-culture of Caco-2 cells and mouse NIH-3T3 fibroblasts [36]. Moreover, the values measured in our system are in line with data from Caco-2 cells in Transwells [37,38] or *in vivo* studies using Ussing chambers [39]. However, it should be noted that TEER values may vary due to different factors such as cell origin and differentiation state, electrode material and positioning, current frequency, medium composition, and experimental setup. Other IoC systems have also showed TEER values that are higher compared to the setup discussed here [25,40,41].

The TEER measurement was more sensitive and reproducible compared to a FITC-dextran permeability assay, which has been described to have poor activity and instable fluorophores [9]. We suggest that TEER measurement is not only more reliable in detecting barrier dynamics and features less variation but further allows to continuously monitor these alterations in real-time, while fluorescence-based permeability assays are considered as end-point measurements [30]. Nevertheless, the presented TEER chip model can also be combined with other biological assays and multiplexing. In this work, it was demonstrated that the tissue can be recovered and used for immunofluorescence staining or downstream biological assays based on cell lysis. This enabled the staining of the tissue to assess the integrity of epithelial and endothelial junctional and adhesion proteins such as E-cadherin, ZO-1, and CD31.

Following studies need to address the demand for multiplexing in OoC systems by integrating and combining additional sensors to measure oxygen levels, pH changes, glucose consumption, and cytokine monitoring complementary to TEER measurements. Within this scope, a previous study was able to demonstrate the measurement of TEER in parallel with oxygen and pH levels in an OoC system [42]. This will pave the way for OoC systems to measure a combination of physical and biochemical parameters in a non-invasive and real-time manner. This is also essential to streamline experiments and to minimize the amount of chips needed for experiments, particularly since most OoC models are still considered low-to mid-throughput [2]. Nonetheless, first studies have already provided high-throughput microplate-based chip platforms with TEER integration that address this limitation [43,44].

It is worth mentioning that membranes with a pore size of 0.4 µm were used in this work. For future studies that want to investigate cell migration between the culture channels, especially of immune cells, pore sizes from 3 to 8 µm would be required. This requires further characterization of the TEER measurement, as membrane properties like material and pore size can have an influence on actual TEER values.

The utilization of the Caco-2 cell line as an epithelial cell surrogate has been used as a reliable cell origin to model the intestinal barrier in a variety of TEER models. Though, it is derived from a tumor origin and might not reflect human-relevant physiological responses, especially regarding transporter expression, metabolic enzyme activity, and drug permeability [45]. Therefore, the results should be validated in an IoC model incorporating adult stem cell-derived epithelial cells [46–48] from patient biopsies in combination with other relevant intestinal cells such as mesenchymal cells, endothelial cells, and immune cells. The use of human induced pluripotent stem cells (hiPSCs) has been a promising avenue for intestinal research since they can be differentiated into relevant cell types originating from one donor [49]. Just recently, a hiPSC-based intestine model was developed, which features self-organization of the epithelial tissue into villus- and crypt-like structures, co-development of myofibroblasts and neurons, and differentiation of the epithelial cells into absorptive and secretory lineages [50]. These advancements will pave the way for IoC models with enhanced physiological relevance to improve clinical translation.

## 5. Conclusion

Collectively, we outlined a biochip platform integrating gold electrodes to measure TEER in real-time by combining it with a novel measurement device facilitating user-friendly plug-and-play handling. Due to the flexibility of the electrode design and chip platform, this approach can be adapted to different OoC models with variable cellular complexities to model biological barriers of interest. While this study provided insights into barrier formation, breakdown, and recovery in an IoC model, future applications include barrier monitoring during microbiota co-culture and infections, inflammatory diseases, and evaluation of gastrointestinal drug toxicities. Expanding the current model with primary cells and incorporating multifunctional sensors will greatly contribute to the development of an IoC platform with non-invasive real-time sensing for personalized medicine and improved clinical translation.

## Data availability statement

All data are included within the article or the supplementary information.

## Acknowledgments

T.K. was funded by the Thüringer Aufbaubank (Germany) – 2021 SD0018 and 2023 SD0029. The authors would like to thank Christian Becker from Dynamic42 GmbH for creating the schematic chip and intestinal model illustrations. We would further like to thank Tobias Voigt for his excellent technical lab support.

## Author contributions

T.K., K.R., S.N. contributed to conceptualization. T.K., N.A., S.B., A.S.M., S.N. contributed to methodology. T.K., S.B. contributed to analysis. T.K., S.B., S.N. contributed to investigation. T.K., S.B., S.N. contributed to writing - original draft preparation. T.K., N.A., S.B., K.R., S.N., contributed to writing - review and editing. K.R., S.N. contributed to supervision. K.R. contributed to funding acquisition.

## References

[1] Zhang B, Korolj A, Lai B F L and Radisic M 2018 Advances in organ-on-a-chip engineering Nat Rev Mater 3 257–78

[2] Low L A, Mummery C, Berridge B R, Austin C P and Tagle D A 2021 Organs-on-chips: into the next decade Nat Rev Drug Discov 20 345–61

[3] Ingber D E 2022 Human organs-on-chips for disease modelling, drug development and personalized medicine Nat Rev Genet 23 467–91

[4] Allaire J M, Crowley S M, Law H T, Chang S-Y, Ko H-J and Vallance B A 2018 The Intestinal Epithelium: Central Coordinator of Mucosal Immunity Trends in Immunology 39 677–96

[5] Pardridge W M 2012 Drug Transport across the Blood–Brain Barrier J Cereb Blood Flow Metab 32 1959–72

[6] Odijk M, Van Der Meer A D, Levner D, Kim H J, Van Der Helm M W, Segerink L I, Frimat J-P, Hamilton G A, Ingber D E and Van Den Berg A 2015 Measuring direct current trans-epithelial electrical resistance in organ-on-a-chip microsystems Lab Chip 15 745–52

[7] Henry O Y F, Villenave R, Cronce M J, Leineweber W D, Benz M A and Ingber D E 2017 Organs-on-chips with integrated electrodes for trans-epithelial electrical resistance (TEER) measurements of human epithelial barrier function Lab Chip 17 2264–71

[8] Yeste J, Illa X, Gutiérrez C, Solé M, Guimerà A and Villa R 2016 Geometric correction factor for transepithelial electrical resistance measurements in transwell and microfluidic cell cultures J. Phys. D: Appl. Phys. 49 375401

[9] Duffy S L and Murphy J T 2001 Colorimetric Assay to Quantify Macromolecule Diffusion across Endothelial Monolayers BioTechniques 31 495–501

[10] Jeon M S, Choi Y Y, Mo S J, Ha J H, Lee Y S, Lee H U, Park S D, Shim J-J, Lee J-L and Chung B G 2022 Contributions of the microbiome to intestinal inflammation in a gut-on-a-chip Nano Converg 9 8

[11] Wang L, Han J, Su W, Li A, Zhang W, Li H, Hu H, Song W, Xu C and Chen J 2023 Gut-on-a-chip for exploring the transport mechanism of Hg(II) Microsyst Nanoeng 9 2

[12] Lucchetti M, Werr G, Johansson S, Barbe L, Grandmougin L, Wilmes P and Tenje M 2024 Integration of multiple flexible electrodes for real-time detection of barrier formation with spatial resolution in a gut-on-chip system Microsyst Nanoeng 10 18

[13] Vera D, García-Díaz M, Torras N, Castillo Ó, Illa X, Villa R, Alvarez M and Martinez E 2024 A 3D bioprinted hydrogel gut-on-chip with integrated electrodes for transepithelial electrical resistance (TEER) measurements Biofabrication 16 035008

[14] Booth R and Kim H 2012 Characterization of a microfluidic in vitro model of the blood-brain barrier (μBBB) Lab Chip 12 1784

[15] Douville N J, Tung Y-C, Li R, Wang J D, El-Sayed M E H and Takayama S 2010 Fabrication of two-layered channel system with embedded electrodes to measure resistance across epithelial and endothelial barriers Anal Chem 82 2505–11

[16] Griep L M, Wolbers F, De Wagenaar B, Ter Braak P M, Weksler B B, Romero I A, Couraud P O, Vermes I, Van Der Meer A D and Van Den Berg A 2013 BBB ON CHIP: microfluidic platform to mechanically and biochemically modulate blood-brain barrier function Biomed Microdevices 15 145–50

[17] Badiola-Mateos M, Di Giuseppe D, Paoli R, Lopez-Martinez M J, Mencattini A, Samitier J and Martinelli E 2021 A novel multi-frequency trans-endothelial electrical resistance (MTEER) sensor array to monitor blood-brain barrier integrity Sensors and Actuators B: Chemical 334 129599

[18] Van Der Helm M W, Odijk M, Frimat J-P, Van Der Meer A D, Eijkel J C T, Van Den Berg A and Segerink L I 2016 Direct quantification of transendothelial electrical resistance in organs-on-chips Biosensors and Bioelectronics 85 924–9

[19] Wei W, Cardes F, Hierlemann A and Modena M M 2023 3D In Vitro Blood-Brain-Barrier Model for Investigating Barrier Insults Advanced Science 10 2205752

[20] Khalid M A U, Kim Y S, Ali M, Lee B G, Cho Y-J and Choi K H 2020 A lung cancer-on-chip platform with integrated biosensors for physiological monitoring and toxicity assessment Biochemical Engineering Journal 155 107469

[21] Holzreuter M A and Segerink L I 2024 Innovative electrode and chip designs for transendothelial electrical resistance measurements in organs-on-chips Lab Chip 24 1121–34

[22] Maurer M, Gresnigt M S, Last A, Wollny T, Berlinghof F, Pospich R, Cseresnyes Z, Medyukhina A, Graf K, Gröger M, Raasch M, Siwczak F, Nietzsche S, Jacobsen I D, Figge M T, Hube B, Huber O and Mosig A S 2019 A three-dimensional immunocompetent intestine-on-chip model as in vitro platform for functional and microbial interaction studies Biomaterials 220 119396

[23] Kaden T, Alonso-Roman R, Akbarimoghaddam P, Mosig A S, Graf K, Raasch M, Hoffmann B, Figge M T, Hube B and Gresnigt M S 2024 Modeling of intravenous caspofungin administration using an intestine-on-chip reveals altered Candida albicans microcolonies and pathogenicity Biomaterials 307 122525

[24] Kaden T, Graf K, Rennert K, Li R, Mosig A S and Raasch M 2023 Evaluation of drug-induced liver toxicity of trovafloxacin and levofloxacin in a human microphysiological liver model Scientific Reports 13 13338

[25] Nazari H, Shrestha J, Naei V Y, Bazaz S R, Sabbagh M, Thiery J P and Warkiani M E 2023 Advances in TEER measurements of biological barriers in microphysiological systems Biosensors and Bioelectronics 234 115355

[26] Shin W and Kim H J 2018 Intestinal barrier dysfunction orchestrates the onset of inflammatory host–microbiome cross-talk in a human gut inflammation-on-a-chip Proc. Natl. Acad. Sci. U.S.A. 115

[27] Zhang L, Li J, Young L H and Caplan M J 2006 AMP-activated protein kinase regulates the assembly of epithelial tight junctions Proc. Natl. Acad. Sci. U.S.A. 103 17272–7

[28] Mendoza C, Nagidi S H, Collett K, Mckell J and Mizrachi D 2022 Calcium regulates the interplay between the tight junction and epithelial adherens junction at the plasma membrane FEBS Letters 596 219–31

[29] Albelda S M, Muller W A, Buck C A and Newman P J 1991 Molecular and cellular properties of PECAM-1 (endoCAM/CD31): a novel vascular cell-cell adhesion molecule. The Journal of cell biology 114 1059–68

[30] Srinivasan B, Kolli A R, Esch M B, Abaci H E, Shuler M L and Hickman J J 2015 TEER Measurement Techniques for In Vitro Barrier Model Systems SLAS Technology 20 107–26

[31] Wang Y I, Abaci H E and Shuler M L 2017 Microfluidic blood-brain barrier model provides in vivo-like barrier properties for drug permeability screening Biotechnol Bioeng 114 184–94

[32] Ramal-Sanchez M, Bravo-Trippetta C, D’Antonio V, Corvaglia E, Kämpfer A A M, Schins R P F, Serafini M and Angelino D 2025 Development and assessment of an intestinal tri-cellular model to investigate the pro/anti-inflammatory potential of digested foods Front. Immunol. 16 1545261

[33] Van Der Helm M W, Odijk M, Frimat J-P, Van Der Meer A D, Eijkel J C T, Van Den Berg A and Segerink L I 2016 Direct quantification of transendothelial electrical resistance in organs-on-chips Biosensors and Bioelectronics 85 924–9

[34] Hidalgo I J, Raub T J and Borchardt R T 1989 Characterization of the human colon carcinoma cell line (Caco-2) as a model system for intestinal epithelial permeability Gastroenterology 96 736–49

[35] Nair A L, Mesch L, Schulz I, Becker H, Raible J, Kiessling H, Werner S, Rothbauer U, Schmees C, Busche M, Trennheuser S, Fricker G and Stelzle M 2021 Parallelizable Microfluidic Platform to Model and Assess In Vitro Cellular Barriers: Technology and Application to Study the Interaction of 3D Tumor Spheroids with Cellular Barriers Biosensors 11 314

[36] Vera D, García-Díaz M, Torras N, Castillo Ó, Illa X, Villa R, Alvarez M and Martinez E 2024 A 3D bioprinted hydrogel gut-on-chip with integrated electrodes for transepithelial electrical resistance (TEER) measurements Biofabrication 16 035008

[37] Béduneau A, Tempesta C, Fimbel S, Pellequer Y, Jannin V, Demarne F and Lamprecht A 2014 A tunable Caco-2/HT29-MTX co-culture model mimicking variable permeabilities of the human intestine obtained by an original seeding procedure European Journal of Pharmaceutics and Biopharmaceutics 87 290–8

[38] Hiebl V, Schachner D, Ladurner A, Heiss E H, Stangl H and Dirsch V M 2020 Caco-2 Cells for Measuring Intestinal Cholesterol Transport - Possibilities and Limitations Biol Proced Online 22 7

[39] Amidon G L, Lee P I and Topp E M 1999 Transport Processes in Pharmaceutical Systems (CRC Press)

[40] Bossink E G B M, Zakharova M, De Bruijn D S, Odijk M and Segerink L I 2021 Measuring barrier function in organ-on-chips with cleanroom-free integration of multiplexable electrodes Lab Chip 21 2040–9

[41] Marrero D, Guimera A, Maes L, Villa R, Alvarez M and Illa X 2023 Organ-on-a-chip with integrated semitransparent organic electrodes for barrier function monitoring Lab Chip 23 1825–34

[42] Izadifar Z, Charrez B, Almeida M, Robben S, Pilobello K, Van Der Graaf-Mas J, Marquez S L, Ferrante T C, Shcherbina K, Gould R, LoGrande N T, Sesay A M and Ingber D E 2024 Organ chips with integrated multifunctional sensors enable continuous metabolic monitoring at controlled oxygen levels Biosensors and Bioelectronics 265 116683

[43] Nicolas A, Schavemaker F, Kosim K, Kurek D, Haarmans M, Bulst M, Lee K, Wegner S, Hankemeier T, Joore J, Domansky K, Lanz H L, Vulto P and Trietsch S J 2021 High throughput transepithelial electrical resistance (TEER) measurements on perfused membrane-free epithelia Lab Chip 21 1676–85

[44] Morelli M, Cabezuelo Rodríguez M and Queiroz K 2024 A high-throughput gut-on-chip platform to study the epithelial responses to enterotoxins Sci Rep 14 5797

[45] Sun D, Lennernas H, Welage L S, Barnett J L, Landowski C P, Foster D, Fleisher D, Lee K-D and Amidon G L 2002 Comparison of human duodenum and Caco-2 gene expression profiles for 12,000 gene sequences tags and correlation with permeability of 26 drugs Pharm Res 19 1400–16

[46] Kasendra M, Tovaglieri A, Sontheimer-Phelps A, Jalili-Firoozinezhad S, Bein A, Chalkiadaki A, Scholl W, Zhang C, Rickner H, Richmond C A, Li H, Breault D T and Ingber D E 2018 Development of a primary human Small Intestine-on-a-Chip using biopsy-derived organoids Sci Rep 8 2871

[47] Beaurivage C, Kanapeckaite A, Loomans C, Erdmann K S, Stallen J and Janssen R A J 2020 Development of a human primary gut-on-a-chip to model inflammatory processes Sci Rep 10 21475

[48] Apostolou A, Panchakshari R A, Banerjee A, Manatakis D V, Paraskevopoulou M D, Luc R, Abu-Ali G, Dimitriou A, Lucchesi C, Kulkarni G, Maulana T I, Kasendra M, Kerns J S, Bleck B, Ewart L, Manolakos E S, Hamilton G A, Giallourakis C and Karalis K 2021 A Novel Microphysiological Colon Platform to Decipher Mechanisms Driving Human Intestinal Permeability Cellular and Molecular Gastroenterology and Hepatology 12 1719–41

[49] Spence J R, Mayhew C N, Rankin S A, Kuhar M F, Vallance J E, Tolle K, Hoskins E E, Kalinichenko V V, Wells S I, Zorn A M, Shroyer N F and Wells J M 2011 Directed differentiation of human pluripotent stem cells into intestinal tissue in vitro Nature 470 105–9

[50] Moerkens R, Mooiweer J, Ramírez-Sánchez A D, Oelen R, Franke L, Wijmenga C, Barrett R J, Jonkers I H and Withoff S 2024 An iPSC-derived small intestine-on-chip with self-organizing epithelial, mesenchymal, and neural cells Cell Reports 43 114247

