## Supplementary Material for "A plug-and-play transepithelial/transendothelial electric resistance (TEER)-upgraded organ-on-chip system to measure barrier dynamics in real-time"

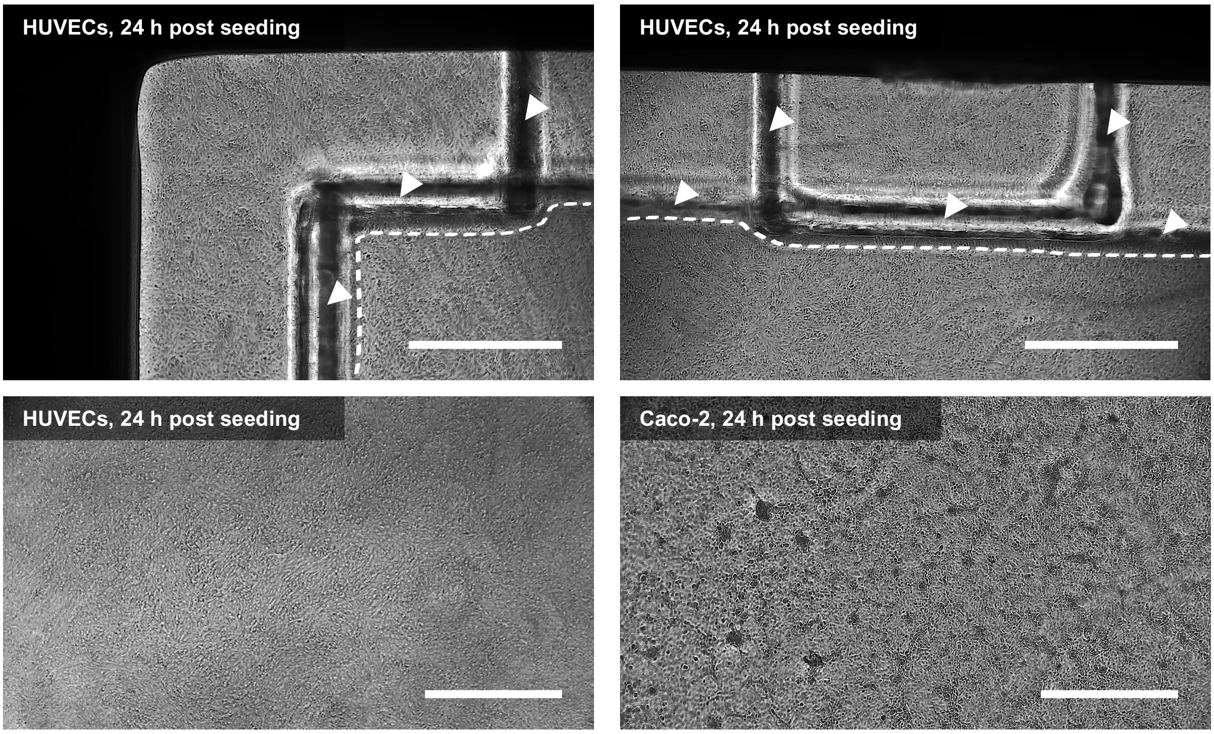


**Supplementary figure 1: Compatibility of integrated TEER electrodes with light microscopic imaging.** Top row images: Representative light microscopic images of HUVECs cultured for 24 h in the TEER-chip. White arrow tips highlight the boarders of the TEER electrodes. Dotted white line outlines the culture area between the electrodes. Bottom row images: Confluent layers of HUVECs (left image) and Caco-2 (right image) on the membrane area observed through the electrodes 24 h post seeding. Scale bars: 1000 µm.

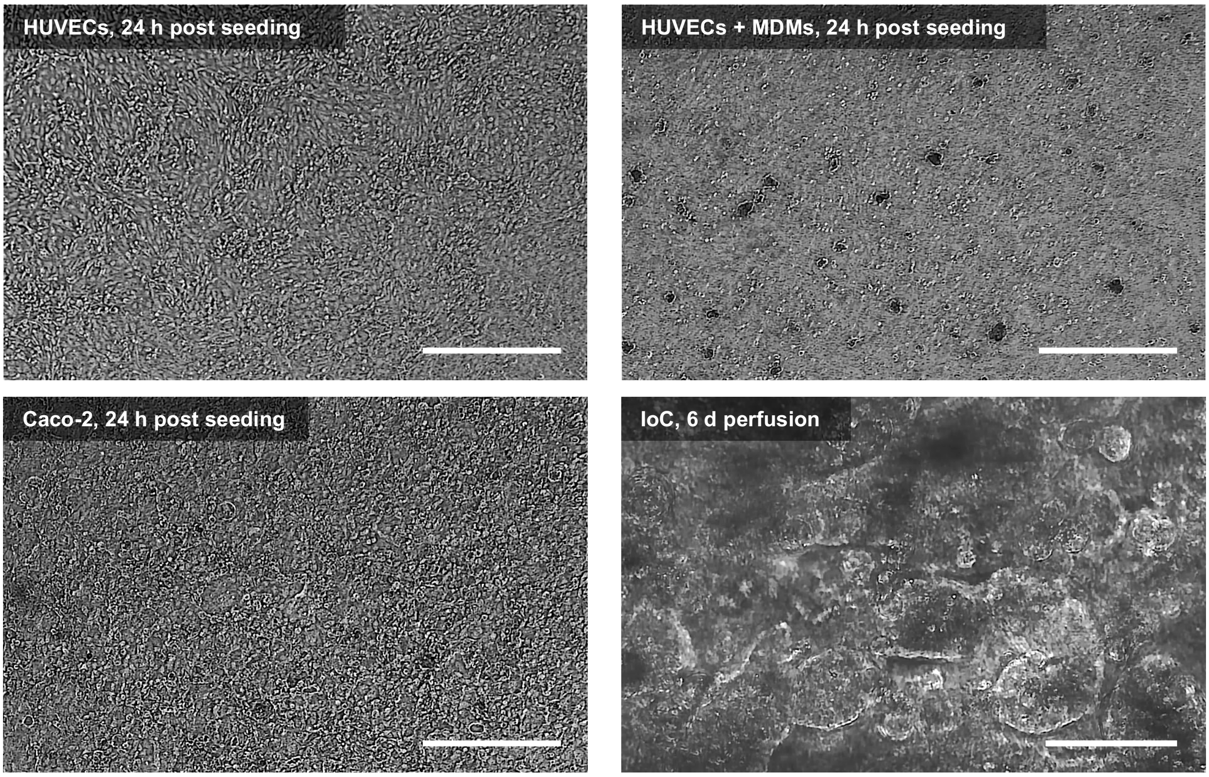


**Supplementary figure 2: Light microscopic monitoring of static IoC assembly and perfusion.** Representative light microscopic images of static IoC assembly with seeding of HUVECs (top left image), MDMs in co-culture with HUVECs (top right image), Caco-2 (bottom left image), and IoC models after six days of perfusion (bottom right image). Scale bars: 500 µm, 100 µm (bottom right image).

**(a)**

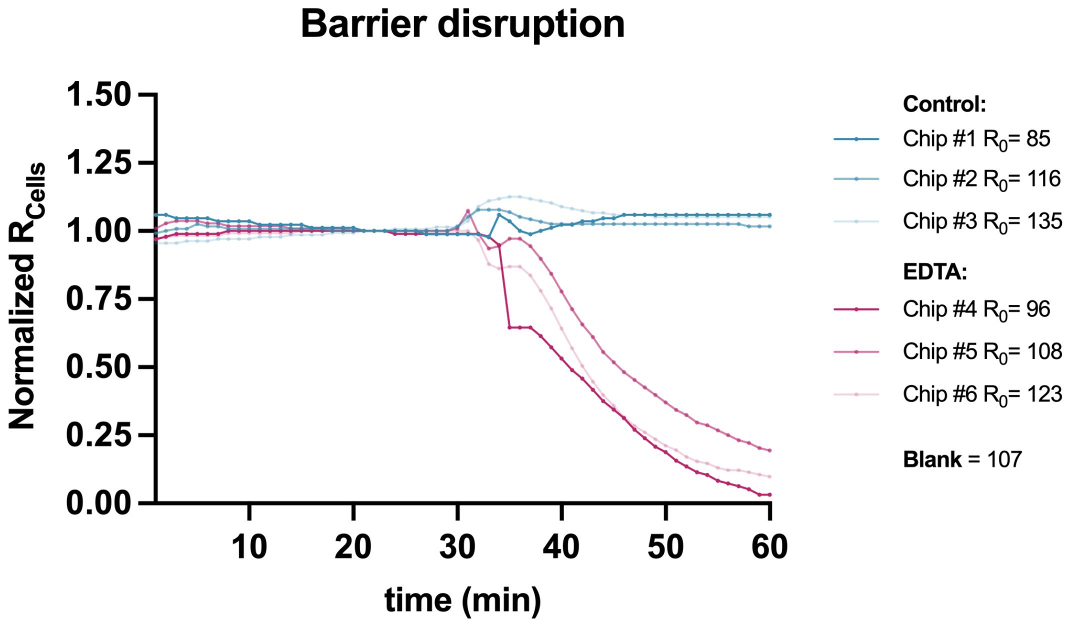


**(b)**

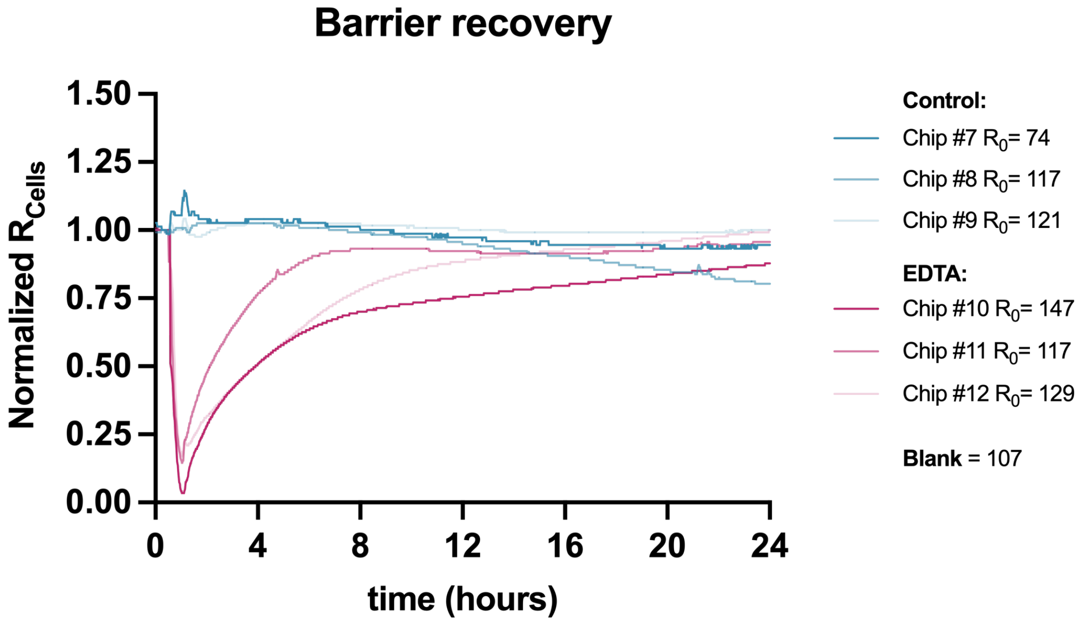


**Supplementary figure 3: Individual chip experiments of barrier disruption and recovery with TEER measurement. (a)** Measurement of TEER during EDTA-induced barrier disruption or **(b)** EDTA-recovery. TEER values were recorded at a measurement interval of 1 min. The minimum blank value was subtracted and values were normalized to the equilibrium TEER value ($R_0$). Each line with or without data points represents an independent chip experiment with control and EDTA in different chip cavities in the same experiment. Results of 3 independent chip experiments (n = 3).
